# Predictors of Population Estimates in a Critically Endangered Species of Black-and-White Colobus Monkeys

**DOI:** 10.1101/2023.08.01.551493

**Authors:** Eva C. Wikberg, Emily Glotfelty, Bright Adu Yeboah, Robert Koranteng, Charles Kodom, Bismark Owusu Anfwi, Afia Boahen

**Author notes:** Corresponding author: Eva Wikberg, Department of Anthropology, University of Texas at San Antonio, One UTSA Circle, San Antonio, TX 78249, USA.

## Abstract

Population monitoring can help us determine population status and trajectory, but it is important to assess what factors may influence the number of individuals counted. In this study we conducted a complete count of the Critically Endangered *Colobus vellerosus* in the forests attached to the Boabeng and Fiema communities in central Ghana. We used 157 repeated counts of the same groups, including both good and unreliable counts to assess what factors predict the number of counted individuals in each group. The number of counted individuals increased with proxies for observation condition, observer experience, and habituation. We therefore recommend observer training and careful planning to increase the chances of having good observation. Then, we used the good counts to calculate the population size and group compositions. The obtained maximum number was 393 individuals in 25 groups. There were no significant differences in group sizes or immature to adult female ratios between groups occupying the older growth forest and groups in other forest types. Although there was still a relatively high immature to adult female ratio indicating that the population size may still increase, it does not appear to grow as rapidly as it used to, based on comparisons with previous population counts. Based on these findings, we recommend priority areas to promote conservation success.

## Introduction

Many primate populations are rapidly declining, and one of the primary drivers of primate population decline is habitat change (Estrada *et al*. 2017). To understand how well populations cope with changing environments and how the population trajectory may be altered by initiatives to mitigate threats, investigators oeen use population monitoring (Nichols and Williams 2006). Population monitoring programs can use a variety of techniques to assess population size and trajectories (Ross and Reeve 2003; Plumptre *et al*. 2013; Campbell *et al*. 2016), and it is important to reflect on the accuracy and precision of the chosen method. One way to assess accuracy is to compare the number generated from the population count with that of study groups with known number of individuals (Kouakou *et al*. 2009). By comparing the actual number of individuals in eight study groups with counts by field assistants not familiar with the study groups during a census of *Colobus vellerosus* (white-thighed or ursine black-and-white colobus) at Boabeng-Fiema Monkey Sancturary in Ghana, Holmes (2011) concluded that the number of individuals counted during the census was approximately 18% lower than the actual number. Also using this methodology in Taï National Park in Côte d’Ivoire, researchers concluded that nest surveys were accurate, because number of *Pan troglodytes* (chimpanzees) estimated from nest data overlapped with the true number (Kouakou *et al*. 2009). Another type of critical analysis focused on evaluating how the number of groups encountered and observed individuals of five primate species may be affected by disturbance when cunng trails to set up the line transects in Salonga National Park, Democratic Republic of the Congo (Bessone *et al*. 2023). When walking the survey route repeatedly on different days, the observers encountered more groups and counted more individuals over time, indicating the primates surveyed were sensitive to this kind of disturbance, and this disturbance had a more prolonged effect on some primate species (Bessone *et al*. 2023). For the more disturbance sensitive species, data from the earlier versus later survey days yielded a three-fold difference in estimated density (Bessone *et al*. 2023). This example illustrates that it is important when designing and interpreting the results of population surveys to be aware of factors that can influence the likelihood of encountering primate groups and the number of individuals observed.

The African colobines are primates with adaptations for an arboreal lifestyle and a diet consisting mostly of leaves and seeds, and the majority of African colobines are threatened by extinction (Wikberg *et al*. 2022). This study focuses on *C. vellerosus* (Fig. 1), which is closely related to *Colobus guereza* (guerezas) and *Colobus polykomos* (western black-and-white colobus) (Oates and Trocco 1983; Ting 2008)*. Colobus vellerosus* is endemic to the Upper Guinean Forest of West Africa, and major threats include habitat change and hunting (McGraw 2005). This led to an elevation of its threat status in 2019 to Critically Endangered, which is the highest threat category on IUCN’s red list before the species goes extinct in the wild (IUCN 2022). There is an estimated 80-87% decline of *C. vellerosus* in Côte d’Ivoire and Ghana with only be 975 mature individuals lee in the wild (IUCN 2022). One of the last remaining large populations of *C. vellerosus* occupies the forests by the villages of Boabeng and Fiema in central Ghana (Wikberg *et al*. 2022). Its population trend is in stark contrast to the trends of most *C. vellerosus* populations throughout its range in West Africa. Repeated complete counts over a 30-year period of the *C. vellerosus* in this area have documented an increased population size from 127 to 451 individuals (Fargey 1992; Saj *et al*. 2005; Wong and Sicotte 2006; Holmes 2011; Kankam and Sicotte 2013; Kankam *et al*. Accepted). The continued presence and size of this population is largely due to the conservation initiatives by people in Boabeng and Fiema who traditionally protected *C. vellerosus* from hunting because of religious taboos (Fargey 1992; Saj *et al*. 2005; Kankam *et al*. 2010). When the taboos eroded over time and only a few dozen monkeys remained in the 1970’s, elders from this community approached Ghana Wildlife Division for governmental protection (Fargey 1992; Saj *et al*. 2005; Kankam *et al*. 2010). In 1990, people from the community initiated the Boabeng-Fiema Monkey Sanctuary (BFMS) ecotourism project for tourists to come and view the monkeys (Fargey 1992; Saj *et al*. 2005; Kankam *et al*. 2010). The population of *C. vellerosus* may also have been increasing rapidly because of local extirpation of local predators. However, threats to this population remain. The closed forest cover in this area has decreased with about 65% (Amankwah *et al*. 2021). Smaller forest fragments and lower tree species richness are associated with lower colobus population densities based on a comparison between 11 forest fragments (Kankam and Sicotte 2013). Forest loss in combination with an increasing population size may lead to increased competition for access to limiting resources (Arseneau-Robar *et al*. 2023; Glotfelty *et al*. In prep.). The increase in home range overlap (Glotfelty 2021), between-group interactions (Arseneau-Robar *et al*. 2023), and within-group interactions (Teichroeb *et al*. 2003; Wikberg *et al*. 2013; Wikberg *et al*. 2014) over time may also lead to increased risk of disease transmission (Nunn and Dokey 2006; MacIntosh *et al*. 2012; Silk *et al*. 2019). Thus, it is important to continue to monitor this population closely to detect changes in size and to assess the accuracy and precision of monitoring methods.

**Figure 1.**
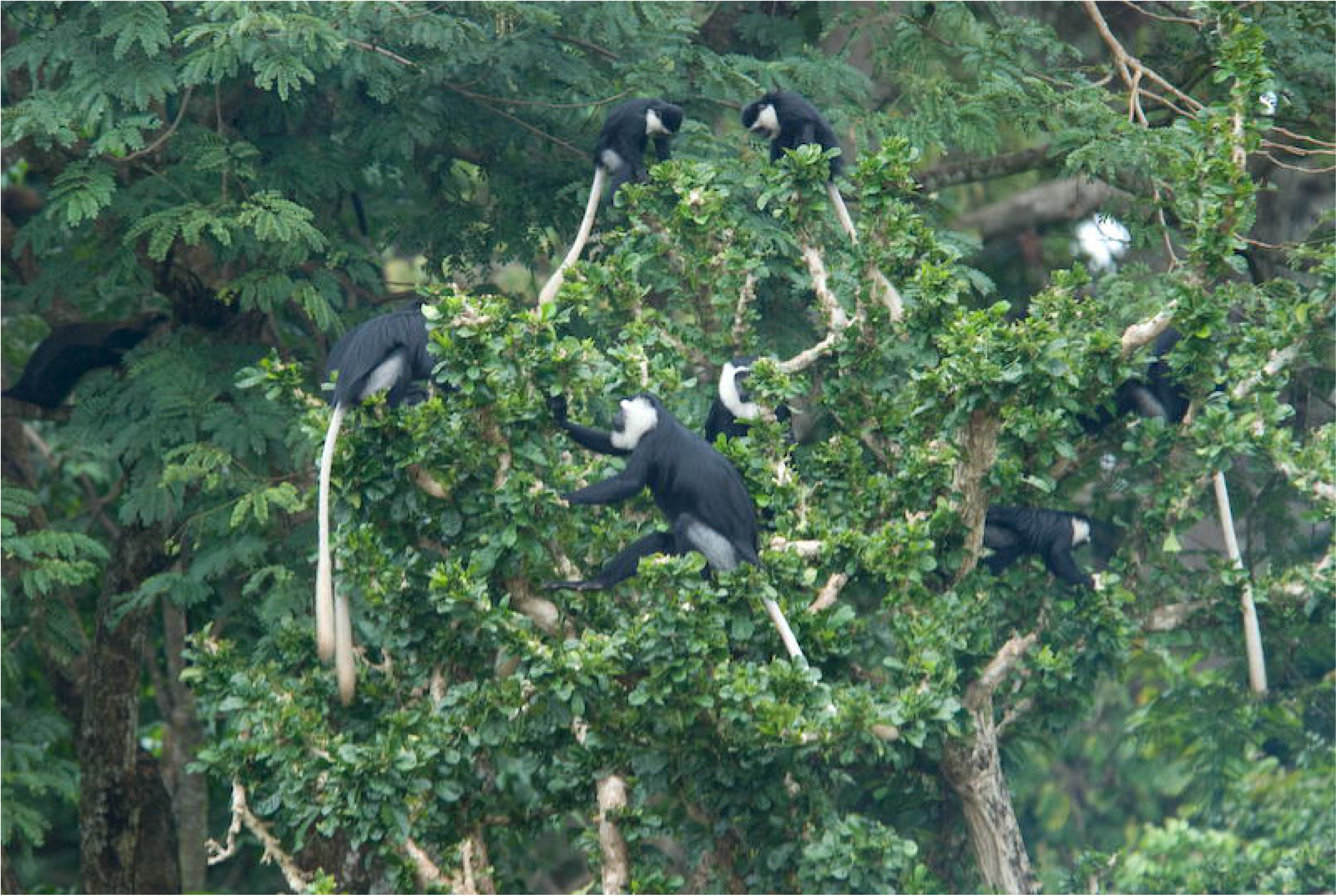
Colobus vellerosus is a diurnal, medium-sized arboreal African colobine who lives in cohesive social groups and has a folivorous diet. Photo credit: Eva C. Wikberg.

Although a previous census concluded that counts were about 18% lower than actual numbers (Holmes 2011), we have not analyzed what factors may improve the chances of obtaining higher counts, more similar to actual numbers. Therefore, the first objective was to determine what factors predicted the observed number of individuals in each encountered group. We predicted that increased habituation level, ideal observation conditions such as high visibility, and observer experience would be associated with a higher number of individuals counted (Ross and Reeve 2003). The second objective was to obtain an updated estimate of the population size and age-sex class composition. We expected: a) a continued positive population trajectory with an estimated population size that would be larger in this study than in previous studies; and b) that the immature to female ratio would be positive and within the range of what has previously been reported from this population, which also would indicate a growing population (Fargey 1992; Saj *et al*. 2005; Wong and Sicotte 2006; Holmes 2011; Kankam and Sicotte 2013; Kankam *et al*. Accepted). The third objective was to evaluate whether group characteristics differed between old growth forest and other forest types, which could possibly indicate that the colobus do better in some forest types than others. We predicted to find larger groups with higher immature to female ratios in the older growth forest based on previous findings of variation in habitat quality and population density across forest fragments (Kankam and Sicotte 2013). Based on our findings, we discuss ways to improve the accuracy of future population counts and conservation action that may aid population growth and expansion.

## Materials and Methods

This study was conducted in a dry semideciduous forest habitat in central Ghana (7°43’N and 1°42’W) (Hall and Swaine 1981). The 1.92 km^2^ land that is set aside for the Boabeng-Fiema Monkey Sanctuary is a mix of primary forest, derived savannah that is regenerating farmland, planted trees, and areas that boarders roads, villages, and farmland (Fargey 1992; Kankam and Sicotte 2013) (Fig. 2). The forest fragment is surrounded by farmland but connected to other, smaller forest fragments via narrow riparian forest corridors monkeys (Fargey 1992; Kankam and Sicotte 2013).

**Figure 2.**
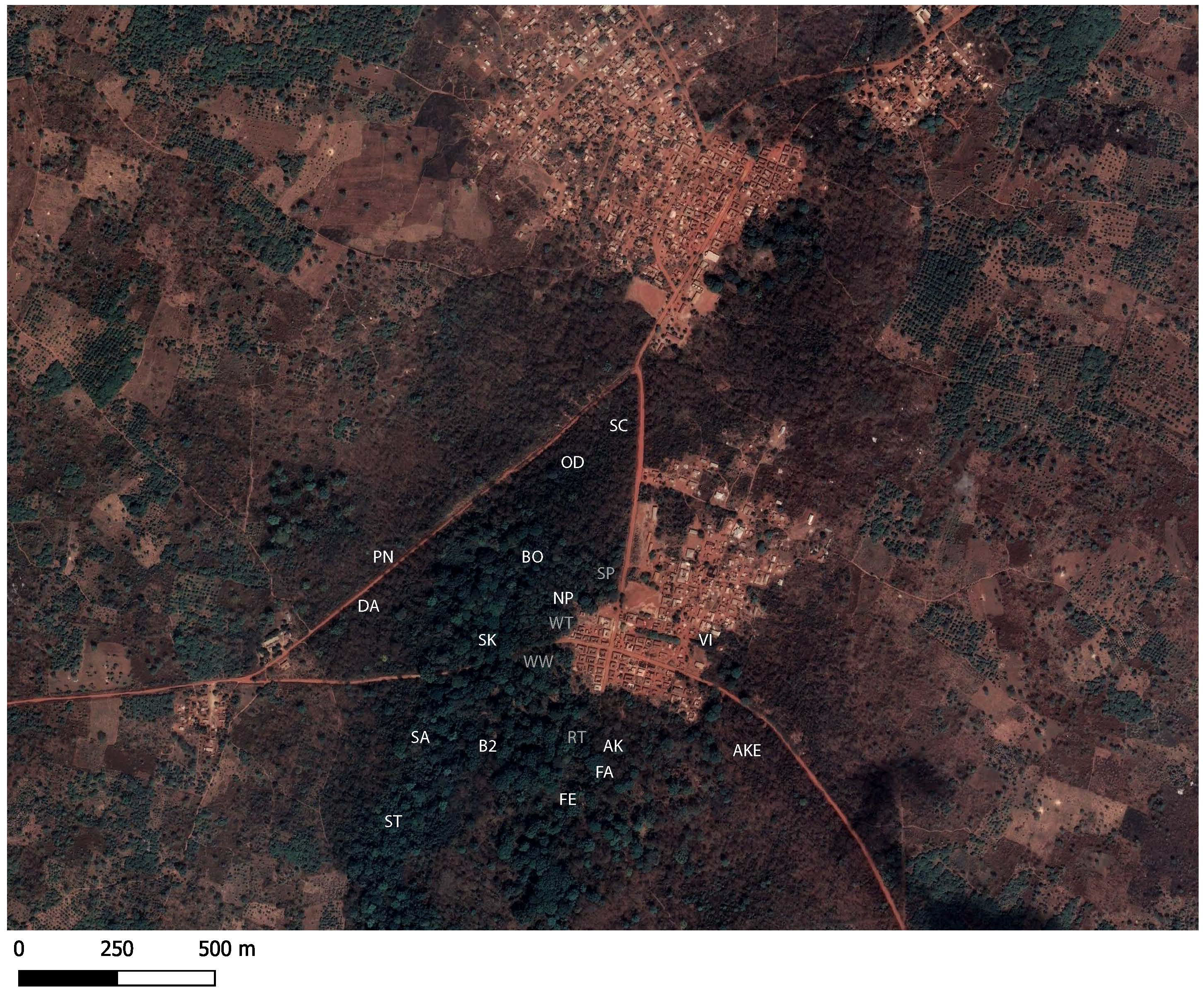
The study area by Boabeng and Fiema in central Ghana with letter codes indicating identities of the groups encountered in the Boabeng forest (grey text = current year-round study group, white text = census group).

*Colobus vellerosus* is a diurnal, medium-sized arboreal African colobine (Saj & Sicotte, 2013; Wikberg *et al*., 2022). They live in cohesive groups with up to 36 individuals (Wong and Sicotte 2006), and groups consist of one or several adult males, one or several adult females, and immatures. As expected for a folivorous primate (Saj and Sicotte 2007; Teichroeb and Sicotte 2009), they spend a high proportion of time resting (Teichroeb *et al*. 2003). All groups in our study area range in proximity to villages, roads, or trails in the forest that are used by community members, tourists, and researchers. Therefore, all groups are at least partly habituated to the presence of humans. Because population counts have been conducted since the 1990’s and field assistants are working in the forest year-round, we have good knowledge of the location of the groups. For groups that are not long-term study groups, group identities are matched up between different census years mostly based on location. Some some group identities are also determined from individuals with unique features (e.g., bent tail, scars, pink nipples, hairless tail) or group-specific behaviors such as one group’s particular agonistic scratch display (i.e., rapidly moving fingers against tree trunks) that have not been observed in any other groups. These characteristics of the species and study population conditions make obtaining a complete count of the population more feasible (Campbell *et al*., 2016; Plumptre *et al*., 2013; Ross & Reeve, 2003). This is the method we used to count the number of individuals in all groups in the Boabeng and Fiema forests, except for the four long-term study groups from which group composition was being recorded three times per week year-round.

We followed the methodology used in some previous population counts at this site (Saj *et al*. 2005; Wong and Sicotte 2006; Holmes 2011). Two to three weeks before the census started, trails were cut in the inaccessible parts of the Boabeng and Fiema forests to be able to find groups that ranged too far away from the existing trails. Six observers conducted the census July 6-30, 2022. Members of our research team were already familiar with how to determine age-sex classes based on their observations of the long-term study groups. During this period, we alternated between group count days (for a total of 10 days) and monitoring sleep trees (for a total of 10 days). On monitoring sleep tree days, all observers walked together starting at 16:00 until one group was found. The most experienced observers identified the group and one less experienced observer stayed with that group until 18:00 when the group would be in their sleep tree while the other observers continued looking for the other groups. They continued like this until each observer was following one group. At 6:00 during group count days, each observer either went to a tree where a group had spent the night (see description of sleep trees below) or they walked trails to find a group. When a group was found, the observer stayed with that group until the group count period ended at 10:00. Once a group was detected, the observer recorded group identity, the location where the group was first found (i.e., a trail, a specific tree, or a location in the village), the number of individuals in each age-sex category that was visible every half hour, whether the count was reliable or not, and comments (i.e., visibility, behaviors, potential errors in age-sex classifications). Individuals were classified as adult male (i.e., large size and continuous white thigh patches), adult female (i.e., smaller than adult males, visible nipples, and white patches on the thighs separated by a thin strip of black fur), juvenile (i.e., moving independently from mother and smaller than adults but larger than infants), infant (i.e., smaller size than juveniles and oeen ventral on adult females), or unknown age-sex class. The observers prioritized walking trails to find groups that had not been contacted yet or from which observers had not yet obtained two good counts. The observers rotated between the groups, and each observer was with a group only once.

We used the repeated counts during the population census days to create a linear mixed model to evaluate whether the observed number of individuals in each encountered group was predicted by variables linked to observation conditions, habituation level, and observer experience. We used the observer’s indication of whether it was a good count as a proxy of observation conditions, as most unreliable counts occur during low visibility. We used time of the day as another proxy of observation conditions, because the colobus groups are typically very cohesive when they are still in their sleep tree while they spread out when they start to move and forage. For the time of the day, we used the minutes since the census start time (6:00). Because we lacked specific indicators of habituation level, we used population identity (Boabeng or Fiema) as a proxy. Groups in Boabeng encounter researchers and tourists more oeen, and we considered the Boabeng groups to have higher habituation level than the Fiema groups. We included census day as a proxy for habituation level and observer experience. Groups that do not typically see observers on a daily basis may become more habituated with each census day and the observers get more experienced in observing and assigning age-sex categories. Observer-specific experience was categorized as 1 if this was the first field season with this study species, 2 if the observer had some past field experience but had not previously conducted a population census, 3 if the observer had conducted a population census and worked as a field assistant in the past, 4 if the observer had previously conducted a population survey and was currently working as a full-time field assistant and, 5 if the observer had conducted multiple population surveys and had been working as a full-time field assistant for about 10 years, and 6 for the observer who had conducted multiple population surveys and worked the longest as a field assistant for the Boabeng-Fiema colobus research project. We performed this analysis in R version 4.1.0 with the packages lme4 glmmTMB (Brooks *et al*. 2017) and multcomp (Hothorn *et al*. 2014), and we used the R packages DHARMa (Hartig, 2021) and performance (Lüdecke *et al*., 2021) to assess model fit and calculate Variance Inflation Factors (VIF). We did not detect any issues with collinearity based on low VIF (range: 1.04-1.10).

We used counts from our four long-term study groups and the census groups for the following calculations. First, we calculated population density as the total number of individuals divided by the area size (1.92km^2^). Second, total biomass (kg/km^2^) was calculated using published weight estimates (males: 8.5 kg, adult females: 6.9 kg, juveniles: 3.85kg (males: 8.5 kg, adult females: 6.9 kg, juveniles: 3.85kg: Oates 1994). For individuals whose age-class could not be determined, we calculated their biomass using the mean weight of the individuals of known age-sex classes. We excluded infants from the biomass calculations to allow for comparisons with previous estimates (Holmes 2011). Finally, the immature to adult female ratio was calculated as the number of infants and juveniles divided by the number of adult females.

We have more long-term ranging data from the Boabeng groups that are study groups or frequently encounter study groups throughout the year. We categorized their home ranges as consisting of older growth forest that typically has multiple canopy layers and over 40% tree cover (e.g., SK in area with dark green tree cover in Fig. 2) or consisting of younger growth forest with a single canopy layer, more shrubs, and herbaceous vegetation (e.g., DA in areas with browner colors in Fig. 2) (Kankam *et al*. 2010; Kankam and Sicotte 2013). We analyzed whether groups in the older versus younger growth forests different in group size or immature to adult female ratio using Mann-Whitney U-tests.

## Results

Each group was contacted two to three times. During each contact day, the observer completed one to eight counts of the group members.

### Count reliability

Of the 178 group counts, 49% were considered good counts, 41% were considered unreliable, and 10% did not have a recorded status. Each group had one to seven good counts. For some good and unreliable counts, the observer described the observation condition. Low visibility when the group was moving in the dense vegetation and/or in the undergrowth was the most common reason cited for unreliable counts (20 comments). Unreliable counts also occurred during intergroup encounters and/or chases (4 comments), when the group was spread out (3 comments), or when it was bad weather (2 comments). Good counts were oeen not accompanied by a comment, but described conditions for good counts included that the group was resting in trees (3 comments) or that they followed a similar path of movement from one large tree to another with high visibility (1 comment). More surprisingly was that some good counts occurred during intergroups and when chases were occurring (3 comments), when there was some movement (2 comments), and when the group was spread out (2 comments).

In a few cases, the observers noted confirmed or possible recording errors. In one case, the observer noted an error in the recorded age-sex class. This was when a jump displaying individual was assumed to be a male but was later confirmed to be a female. Males are jump displaying more oeen than females, and it is also difficult to sex the individuals when they are moving fast in the canopy. In another case, the observer could hear an infant squealing but was not able to see it, and not detecting infants would deflate group size. The observers also noted possible errors in their recordings that would have inflated the group size count. Two of these cases occurred during intergroup encounters. In the third case, the observer noted that they may have double-counted one individual.

### Predictors of observed group size

In the analysis of observed group size, we included the 158 group counts from 21 groups when at least some individuals were visible and the observers had recorded whether their count was reliable. The observed number of individuals was lower when the count was classified as unreliable by the observer (estimate: -3.43, 95% CI: -4.27 – -2.58), increased with observer experience level (estimate: 1.15, 95% CI: 0.59 – 1.69), and number of days since the census started on July 7 (estimate: 0.78, 95% CI: 0.22 – 1.35) (Fig. 3). The observed number of individuals was not predicted by hours since the start of the census at 6:00 in the morning (estimate: -0.3, 95% CI: -0.71 – 0.11) or by population ID (estimate: -0.56, 95% CI: - 4.04 – 2.92) (Fig. 3). The model explained 23% of the observed variation without random effects and 73% of the observed variation with random effects.

**Figure 3.**
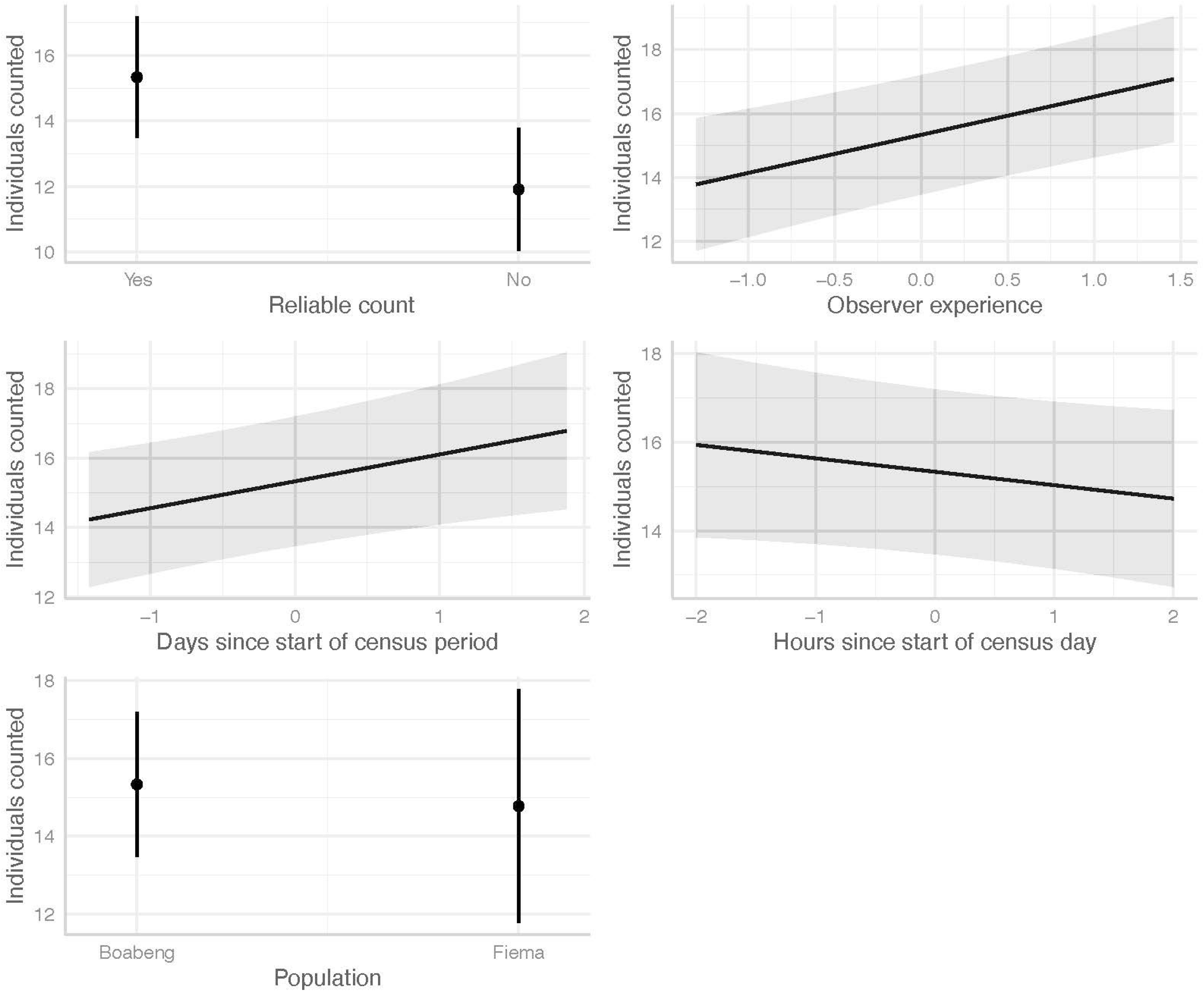
The predicted relationship between the number of individuals counted in each social group and whether the observer classified it as a reliable count, observer experience, days since the first census day, hours since start of the census that day, and whether the group ranged by Boabeng or Fiema. Shaded areas represent the 95% confidence interval, and the y-axis with numerical values has been square root transformed.

### Population size and group composition

Observers counted 19 groups in the Boabeng forest and 6 groups in the Fiema forest (Table 1). One Boabeng group and two Fiema groups that had been present during previous population counts could not be located. Only including the counts classified as good counts by the observer, the number of counted individuals ranged from a minimum of 349 to a maximum of 381. The calculated density ranged from 181.77 to 198.44 individuals/km^2^ if using minimum versus maximum counts. If using the maximum number of individuals counted, the calculated biomass was 663.00 kg for adult males, 938.40 kg for adult females, 350.35 kg for juveniles, and 127.98 kg for individuals of unknown age-sex class. If including individuals with known and unknown age-sex classes, the total biomass is 2,079.73 kg.

**Table 1.**
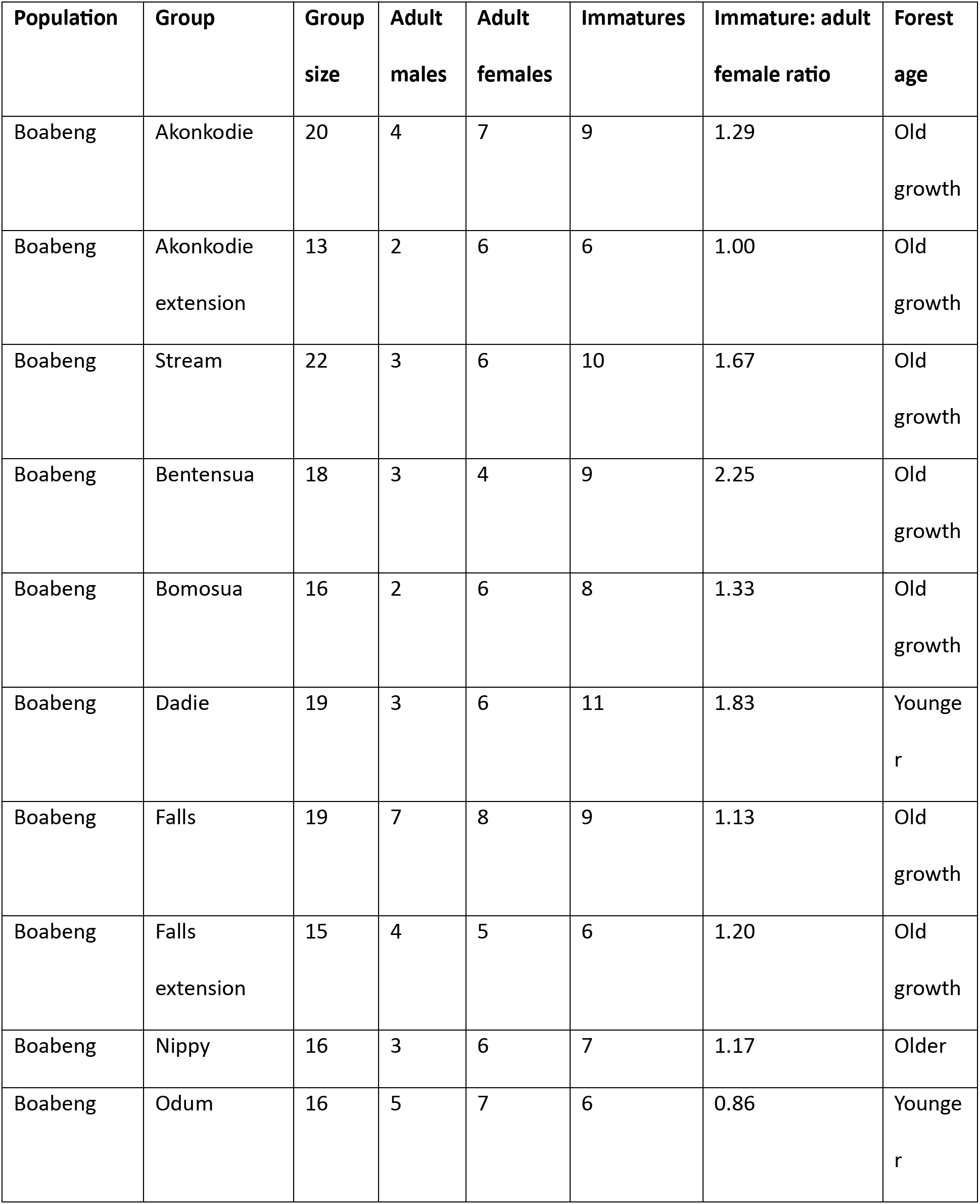
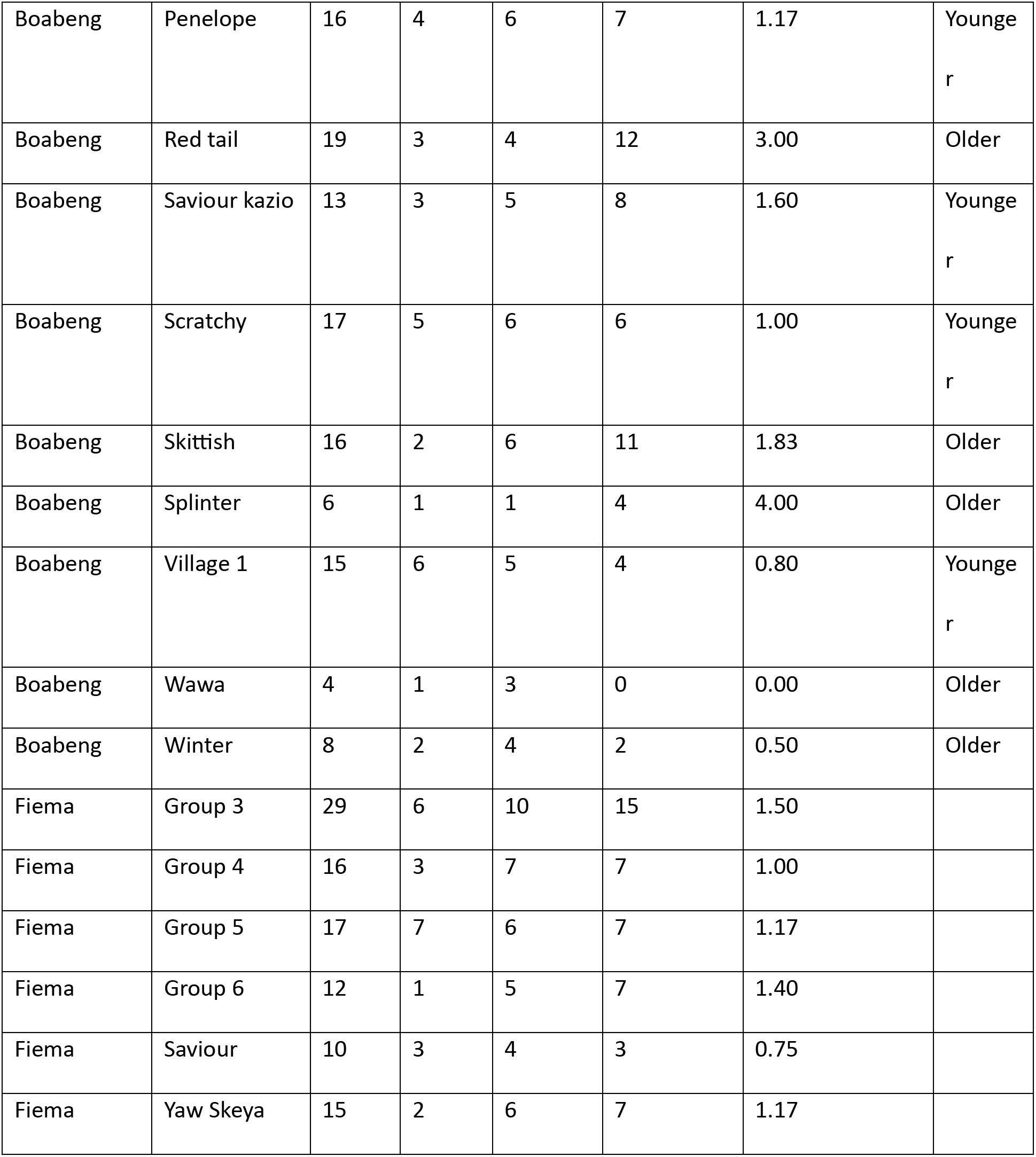
Composition of the groups inhabiting the forests by Boabeng and Fiema.

Each group contained 1 to 7 adult males, 1 to 10 adult females, and 0 to 15 immatures based on the maximum numbers observed in each group (Table 1). The median group size was 16 both in the older growth forest (range: 4 to 22) and in the younger growth forest (range: 13 to 19), and there was no significant difference in size between groups in these two forest types (Mann-Whitney U-test, W = 39, p = 1.10, N = 19). We counted a total of 136 adult females and 178 immatures (using maximum values for number of individuals counted), which yields an overall immature to adult female ratio of 1.31. Although the median immature to adult female ratio was slightly higher in groups occupying the older growth (1.20, range: 0 to 4) than that in groups in the younger forest (1.08, range: 0.80 to 1.83), there was no significant difference in immature to adult female ratio between groups in these two forest types (Mann-Whitney U-test, W = 49.5, p = 0.38, N = 19).

## Discussion

### Predictors of observed group size

We conducted a complete count of the Boabeng-Fiema population of the Critically Endangered *Colobus vellerosus*. We used repeated counts of the same groups to analyze what factors predicted the number of counted individuals in each group. The observed number of individuals was predicted by whether the count was classified as reliable by the observer. Unreliable counts oeen occurred during low visibility as expected (Ross and Reeve 2003). However, observers were still able to obtain good counts during some challenging observation conditions like during intergroup encounters. Likely due to their prior knowledge of colobus behaviors, they could in some cases single out extra-group individuals even when they were attached to the group being counted. Observer expertise is likely very important for the accuracy and precision of complete counts (Ross and Reeve 2003). As expected, the observed number of individuals increased with observer experience level. The observed number of individuals also increased with number of days since the census started, which could be another proxy for observer experience with the underlying reasoning that observers get more experienced in observing and assigning age-sex categories during the study. However, it can also be a proxy for increased habituation level of the groups over time, making it easier to count individuals from groups that do not typically see observers on a regular basis. In contrast, Bessone and collegues (2023) argue that an increased number of individuals counted over time was due to decreased disturbance with increasing time since trails were cut. We believe it is unlikely that trail cunng would have affected the monkeys’ behaviors in our study as the trails were cut two to three weeks before the start of the census and all individuals are at least partly habituated to humans. In contrast to our predictions, there was no significant difference in observed number of individuals between the two populations. We had expected it to be more difficult to count individuals in the Fiema forest as these groups encounter tourists and researchers less frequently than the groups in the Boabeng forest do. Although our results could indicate that population ID is a poor proxy for habituation and/or that the habituation level of groups in Boabeng and Fiema are similar, the Fiema groups directed more display to and fled from the only foreign researcher on the census team. Similar behaviors were reported by Holmes (2011), who also described how some colobus groups would flee at the sight of foreign objects such as binoculars. However, she was still able to obtain reliable counts when the group members leaped between trees or fled into taller trees with high visibility (Holmes, 2011). Finally, the study species activity patterns may affect the likelihood of obtaining complete counts (Ross and Reeve 2003). We predicted that the observed number of individuals would decrease with the number of hours since the start of the census day, because the groups typically spend the night in tall trees (Teichroeb *et al*. 2012) and are more cohesive and visible early in the morning before they start to travel and forage in the dense undergrowth closer to the ground. In contrast to our prediction, time of the day did not predict the number of observed individuals, and we conclude that variables linked to observation conditions, observer experience, and habituation are more important.

Based on these findings, we provide the following recommendations to improve the accuracy of future population estimates. Observer experience was an important predictor of number of individuals counted, and ideally, people on the census team should be well-trained before the census starts. For future census work, it may also be worthwhile to have a team of observers locating all non-study groups before the census starts to become more familiar with their ranging pattern and increase habituation levels. Our descriptive data suggest that the less habituated monkeys responded differently to the foreign researcher on our census team, which should be considered when deciding who will focus on which forest fragment. Because the observers’ perceived quality of the count was an important predictor of number of individuals observed, we recommend that they keep detailed notes on the perceived quality of their count, conditions that may affect the observed number, and any uncertainties about double-counting, mis-sexing individuals, or including extra-group individuals in their count.

Finally, we conducted our population census during the rainy season due to time constraints, similar to previous counts of this population (Saj *et al*. 2005; Wong and Sicotte 2006; Kankam *et al*. 2010; Holmes 2011; Kankam and Sicotte 2013). However, it is ideal to perform the complete counts during seasons when it is easier to count all individuals (Ross and Reeve 2003). There is less foliage during the dry season, and it would be easier to spot individuals when the visibility is higher. Thus, there may be several ways to refine population monitoring methods, which may make it possible detect smaller changes in population trajectories.

### Population size and group composition

During our complete count of *C. vellerosus* in the summer of 2022, observers contacted 25 groups and counted 350 - 393 individuals in the area by Boabeng and Fiema. The number of mature individuals in this population represent approximately 20% of the estimated 975 mature *C. vellerosus* living in the wild (Matsuda Goodwin *et al*. 2020), and our study population is largest known population of *C. vellerosus* (Wikberg *et al*. 2022). Based on the maximum observed number of individuals, the population density is estimated to be 198.44 individuals/km^2^. Several populations of red colobus (*Piliocolobus spp.*), which is a closely related taxon to the black-and-white colobus, also occur at similar or higher population densities (Wikberg *et al*. 2022). The density of our study population is at the upper range reported from other populations of black-and-white colobus monkeys (Wikberg *et al*. 2022). Notably, the *C. vellerosus* population density in Dinaoudi sacred grove in Côte d’Ivoire was a staggering 1000 individuals/km^2^, but it only consisted of 30 individuals in a 3 ha fragment (Gonedelé Bi *et al*. 2010). Thus, while some colobine population can reach high densities, several of these cases likely consist of populations being compressed in shrinking forest patches. It is uncertain whether such compressed populations can persist long-term, especially if the population size is small.

The maximum number of counted individuals during our 2022 census was 393, which is 58 individuals less than what was counted during Kankam and colleagues’ (Accepted) 2020 census. However, Kankam and colleagues’ (Accepted) worked in teams of two, and it is possible that this increases the number of observed individuals. Our methodology followed that of Holmes’ (2011) who determined that this method underestimates the size with approximately 18% (Holmes 2011). If this is the case, the actual number of individuals in the population in 2022 would be 463 individuals, which is similar to the 451 individuals counted during Kankam and colleagues’ (Accepted) census. Both complete count methods used at this site should lead to an accurate number of groups, because we have a good understanding of the number and approximate locations of groups. Three known groups were missing during our census, and our total number of groups was two groups fewer than during Kankam and colleagues’ (Accepted) 2020 census. These groups may have dissolved or moved out of the study area. Groups have previously been reported not to reside year-round in Fiema (Kankam *et al*., 2010).

When comparing our population count to those conducted in the past, the population appears to have increased at an average rate of 6.75 individuals per year based on the maximum number during the current census in 2022 (393 individuals) and the maximum number counted in 2010 (312 individuals), which was the last census that used the exact same methodology (Holmes 2011). This is only about half the rate at which the population size increased between the census years 2003, 2007, and 2010 (Wong and Sicotte 2006; Kankam *et al*. 2010; Holmes 2011).

On a positive note, the overall immature to adult female ratio during our population count was at the high end of those reported previously (Saj *et al*. 2005; Wong and Sicotte 2006; Holmes 2011; Kankam and Sicotte 2013; Kankam *et al*. Accepted), and positive numbers suggest that the population is still on a positive population trajectory. In contrast, having more adult females than infants and juveniles would likely signal a decreasing population trend as the case was in the decreasing *Alou<a palliata* (mantled howler) populations in Panama and Costa Rica (Heitne *et al*. 1976). Also, we did not find any significant differences in group size or immature to adult female ratios between groups inhabiting the old growth forest versus younger forest. Thus, the colobus may be able to survive and reproduce well even in forests that consist of regenerating farmland and cultivated trees. Cultivated tree species make up (during some months) 32% of the diet of *Aloua<a guariba clamitans* (brown howlers) in Rio Grande do Sul State, Brazil (Chaves and Bicca-Marques 2017). The authors conclude that cultivated tree species that both humans and non-human primates utilize can have an important conservation value (Chaves and Bicca-Marques 2017), and this may be especially true in human-dominated landscapes.

### Evidence-based conservation

It is important to keep in mind that determining actual population trajectories can be difficult due to between-year variation and sampling effects (Nichols and Williams 2006), and it may take up 10 years of data to accurately determine population trends (Maxwell and Jennings 2005). This timeframe would be too long to wait before incorporating the results of studies in management plans (Nichols and Williams 2006), because the threats to primate survival are escalating rapidly (Estrada *et al*. 2017). For example, a small population of *C. vellerosus* in Soko sacred grove in Côte d’Ivoire went locally extinct between surveys conducted three years apart (Gonedelé Bi *et al*. 2010). Because the population of *C. vellerosus* in the Boabeng and Fiema forests appears to be increasing less rapidly than two decades ago, it is important to evaluate why this may be, what actions should be put in place to promote future population growth and expansion, and then follow best practices to assess the effectiveness of these actions (Junker *et al*. 2020; Christie *et al*. 2021).

Our study species *C. vellerosus* and some other species of African colobines show a greater degree of behavioral flexibility than anticipated for species adapted to a highly specialized diet, and they can persist in human-modified environments (Wikberg *et al*. 2022). The ability to use human-modified landscapes such as human settlements and secondary forests are associated with reduced extinction risk (Galán-Acedo *et al*. 2019). Less strictly arboreal species with increased dietary diversity are more likely to use human-modified environments (Galán-Acedo *et al*. 2019), and dietary diversity also reduces extinction risk (Jernvall and Wright 1998; Machado *et al*. 2023). This behavioral flexibility may increase their chances of population persistence in changing environments (Buskirk, 2012; Beever *et al*. 2017). However, there may be a limit to how much the individuals can change their diet and habitat use. Similarity, howler monkeys also use individual behavioral flexibility to cope with habitat loss, but they seem less likely to persist long-term in smaller compared to larger habitat patches (Bicca-Marques *et al*. 2020). Indeed, behavioral changes may not always be adaptive or sufficient to cope with environmental changes, and these changes may indicate a future population collapse (Berger-Tal *et al*. 2011).

Alternatively, it may be that the forest cannot accommodate a higher number of colobus monkeys. In line with this notion, we have observed behavioral changes in our study groups over time that indicate increased competition for food resources (Wikberg *et al*. 2013; Arseneau-Robar *et al*. 2023; Glotfelty *et al*. In prep.). Under this scenario, more individuals may be motivated to disperse either temporarily or permanently to the surrounding forest fragments (Jones 2005). Indeed, seven nearby forest fragments have been recolonized by colobus in the last decades years, most likely by dispersing individuals from the Boabeng and Fiema fragments (Wong and Sicotte 2006; Kankam *et al*. 2010).

Although *C. vellerosus* at our study site has recolonized some of the surrounding forest fragments, some fragments remain unoccupied or only contain a very small number of individuals (Wong and Sicotte 2006; Holmes 2011; Kankam and Sicotte 2013; Kankam *et al*. Accepted). Planting trees and building landscapes that take the needs of forest-dwelling animals and humans into account could facilitate dispersal between fragments (Arroyo-Rodríguez *et al*. 2020). To promote dispersal of *Leontopithecus chrysopygus* (black lion tamarin) in the Pontal do Paranapanema region of São Paulo State, Brazil, local community members and other stakeholders built forest corridors and stepping stones (Chazdon *et al*. 2020). It is important to evaluate whether the location of these corridors and steppingstones compete with socioeconomic and cultural values for that land (Ruiz-López *et al*. 2022). For example, planting a series of steppingstones using habitat not suitable for farmland may not interfere with human livelihoods but could still link surrounding fragments to the Kibale National Park in Uganda and encourage *Piliocolobus tephrosceles* (red colobus) dispersal (Ruiz-López *et al*. 2022). It is possible that similar approaches to building landscapes that considers the needs of the colobus monkeys and humans could facilitate dispersal between forest fragments at our study site.

In addition to facilitating movement between fragments, it is also important to improve chances for population persistence in the fragments once a population is established there. The colobus monkeys are arboreal and leaves from large trees make up the majority of the colobus diet (Saj and Sicotte 2007; Teichroeb and Sicotte 2009). Therefore, the rapid loss of forest in this area is concerning (Kankam *et al*. 2010; Amankwah *et al*. 2021). Some of the forest loss has been accidental due to forest fires. Although our study group started using a burnt area relatively soon aeer the regrowth of vegetation (CK, personal observation), severe fires can have a long-term effect on primates. *Symphalangus syndactylus* (siamangs) did not incorporate heavily burnt areas while they did resume ranging in other areas with less severe fire effects within 18 years aeer a forest fire (Lappan *et al*. 2021). Improved forest protection could be achieved by preventing additional cunng of large trees and allocating more resources to fire management. It may also be possible to promote further increase and/or persistence of the *C. vellerosus* population by planting trees in certain areas with low tree coverage to increase habitat carrying capacity. Unfortunately, this is a slow process that requires long-term care to prevent the trees being outcompeted by faster growing plants.

The Boabeng and Fiema communities have also taken action by sharing revenue from the ecotourism projects with the surrounding communities, which is likely an important incentive for them to protect the colobus as they do not have the same traditional beliefs as the Boabeng and Fiema communities do (Kankam *et al*. 2010). Revenue sharing could increase the chances for population expansion and persistence. By finding ways to mitigate threats to non-human primates while also considering human livelihoods and improving human well-being, we may be able to enhance the conservation outlooks for the many threatened non-human primates living in close proximity to humans (Kareiva and Marvier 2012).

## Ethics Statement

Our research adheres to ABS/ASAB guidelines, IPS Code of Best Practices for Field Primatology, and the laws of Ghana. Data collection was approved by the Boabeng-Fiema Monkey Sanctuary’s management committee, the Ghana Wildlife Division, and the University of Texas at San Antonio Institutional Animal Care and Use Committee (CO001-05-25).

## Acknowledgement

We would like to thank the Boabeng-Fiema Monkey Sanctuary management committee and the Ghana Wildlife Division for permission to conduct this research and the community members for providing additional information of where to find colobus groups. We thank the Leakey Foundation (ECW, BAY), the International Primatological Society (EG), and the University of Texas at San Antonio (ECW, BAY, EG) for funding.

